# Exposure to unpredictable trips and slips while walking can improve balance recovery responses with minimum predictive gait alterations

**DOI:** 10.1101/333989

**Authors:** Yoshiro Okubo, Matthew A Brodie, Daina L Sturnieks, Cameron Hicks, Hilary Carter, Barbara Toson, Stephen R Lord

## Abstract

**INTRODUCTION:** This study aimed to determine if repeated exposure to unpredictable trips and slips while walking can improve balance recovery responses when predictive gait alterations (e.g. slowing down) are minimised.

**METHODS:** Ten young adults walked on a 10-m walkway that induced slips and trips in fixed and random locations. Participants were exposed to a total of 12 slips, 12 trips and 6 non-perturbed walks in three conditions: 1) right leg fixed location, 2) left leg fixed location and 3) random leg and location. Kinematics during non-perturbed walks and previous and recovery steps were analysed.

**RESULTS:** Throughout the three conditions, participants walked with similar gait speed, step length and cadence(*p*>0.05). Participants’ extrapolated centre of mass (XCoM) was anteriorly shifted immediately before slips at the fixed location (*p*<0.01), but this predictive gait alteration did not transfer to random perturbation locations. Improved balance recovery from trips in the random location was indicated by increased margin of stability and step length during recovery steps (*p*<0.05). Changes in balance recovery from slips in the random location was shown by reduced backward XCoM displacement and reduced slip speed during recovery steps (*p*<0.05).

**CONCLUSIONS:** Even in the absence of most predictive gait alterations, balance recovery responses to trips and slips were improved through exposure to repeated unpredictable perturbations. A common predictive gait alteration to lean forward immediately before a slip was not useful when the perturbation location was unpredictable. Training balance recovery with unpredictable perturbations may be beneficial to fall avoidance in everyday life.

## Introduction

Falls in older people are a major cause of fractures, physical dependency, mortality and economic burden [1]. Perturbation training is an emerging paradigm that utilizes repeated exposure to sudden external perturbations, aimed at inducing locomotor adaptations and training balance recovery responses specifically required for fall avoidance [2, 3]. Theoretically, exposure to postural perturbations in a safe environment enables training of balance recovery responses that are critical for avoiding falls due to unexpected slips and trips in everyday life.

There is good evidence that both young and older adults can adapt their gait and improve their balance responses in response to repeated perturbations and substantially reduce falls in the laboratory within a few trials. However, if not regulated, participants tend to shorten their steps and shift their centre of mass [CoM]) to minimize the disturbing effects of the known upcoming perturbations [5, 11]. Although it has been reported that these predictive gait alterations are transferable across different conditions [11–13], many studies have used repeated perturbations at a consistent location which could make later trials highly predictable. Moreover, many studies used a single perturbation type (e.g. slips only) which could result in an over-adaptation and increased vulnerability to opposing perturbation types (e.g. trips). Therefore, it is questionable if such a gait strategy against predictable perturbations acquired in a laboratory setting is useful for situations where perturbations occur unexpectedly as in everyday life.

Moreover, although many studies have reported significant improvements in reactive balance control following repeated perturbations [5–7, 9, 14–16], a recent systematic review found that 80% of the included studies did not control for the effect of predictive gait alterations (e.g. including sufficient washout walks) [4]. Anticipation of upcoming perturbations and related changes in the approach walk (slower gait speed, shorter step lengths and anticipatory postural adjustments) can diminish the perturbation magnitude and therefore possibly overestimate the improvements in reactive balance control reported [4]. Thus, the evaluation of balance recovery responses during walking has proved to be a significant challenge because unpredictability and magnitude of postural perturbations are difficult to maintain over repeated trials

To address the above issues, we developed a 10m perturbation walkway that can generate both trips and slips (i.e. opposing type perturbations) in varied locations while regulating step length, cadence and gait speed. This system, therefore, minimises predictive gait alterations during repeated perturbations, maintaining a greater level of unpredictability, as is in real life. The primary objective of this study was to test the hypothesis: Exposure to unpredictable perturbations can improve balance recovery responses in the absence of predictive gait alterations. In addition, to evaluate the importance of minimizing predictive gait alterations, we also tested following secondary hypothesis: Predictive gait alterations (e.g. shift of CoM) acquired through exposure to perturbations at a fixed condition would not transfer to conditions where both the type and location of perturbation were randomly presented and highly unpredictable. Findings from this study may contribute significantly to the design of effective exercise interventions using perturbations for fall prevention.

## Methods

### Participants

Eleven healthy adults aged 20-40 years provided written informed consent prior to participating in this study. Inclusion criteria were: aged between 20-40 years, living independently, and able to stand or walk for 20 minutes unassisted. Exclusion criteria were diagnosis of osteoporosis or neurological impairment that restrict activities of daily living (none was excluded). The study protocol was approved by the University of New South Wales Human Research Ethics Committee (HC16227).

### Apparatus

The trip and slip perturbation system was built into a 10m walkway consisting of wooden decking with stepping tiles atop (Fig 1). A slip was induced by a movable tile on two hidden low-friction rails with linear bearings that could slide forward up to 70cm upon foot contact. The slipping tile was locked or unlocked as appropriate using a concealed wedge. A trip was induced using a 14cm height tripping board that could spring up from the walkway. The trip board comprised two wooden walkway boards (14cm combined height) – a trip height consistent with previous studies that used 11cm to 15cm high obstacles [16–19]. A trained tester triggered the tripping board with a wireless controller at mid-swing of the gait cycle [17]. The tripping board and slipping tile were not visually detectable and could be moved to any location along the walkway. The participants faced away from the walkway before each trial so that the positioning of the hazards could not be seen.

**Fig 1.**
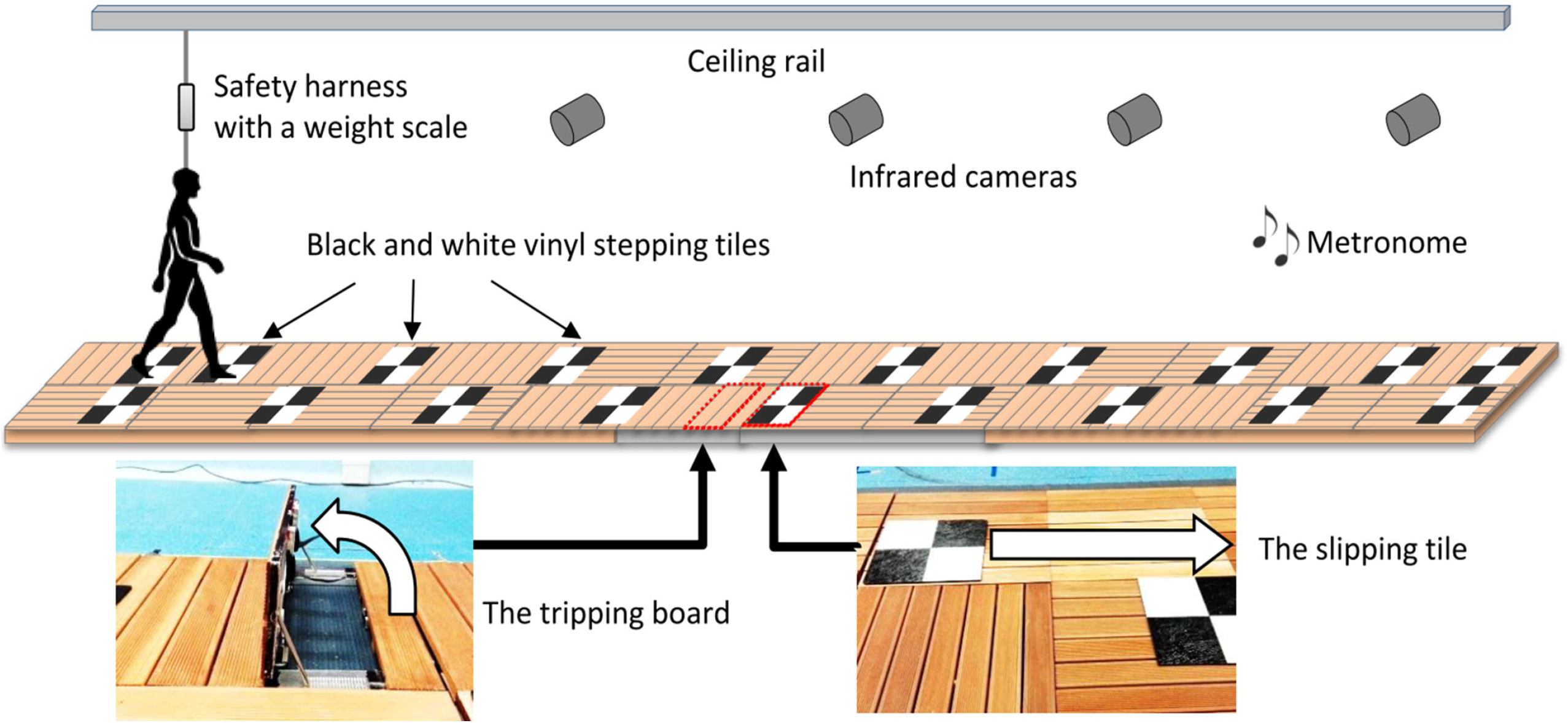
The trip and slip perturbation system used for this study. A slip was induced by a movable tile on two hidden low-friction rails with linear bearings that could be unlocked to slide up to 70cm upon foot contact. A trip was induced using a 14cm height tripping board that could be triggered to spring up from the walkway. The tripping board and slipping tile were not visually detectable and could be moved to any location along the walkway. Black and white vinyl stepping tiles were placed on the walkway to reproduce individual’s step length and a metronome was set to the individual’s cadence.

### Experimental protocol

Initially, participants walked at usual pace for 3 lengths of a 5.7m (active area: 4.8m) electronic walkway (GAITRite ® mat, v4.0, 2010 CIR Systems, USA) to calculate average step length and cadence. A one-meter approach was provided so that participants were walking at their normal pace when on the walkway. Participants were also instructed to continue walking for one meter beyond the walkway to ensure that the walking pace was kept consistent throughout the task. Using this information, black and white vinyl stepping tiles were placed on the perturbation walkway to regulate each individual’s usual step length and a metronome was set to regulate their usual cadence (Fig 1).

Participants were secured with a ceiling-mounted full body harness to avoid contact with the ground if they fell following a perturbation. All participants were exposed to slips and trips in 3 sessions of different conditions (Table 1). In each session, 4 slips, 4 trips and 2 non-perturbed walk (catch) trials were administered. To minimize predictive gait alterations, slips and trips were mixed in each condition. In condition 1, the hazards were placed on the right side of the walkway at a fixed middle location. In condition 2 the hazards were placed on the left side of the walkway at a fixed middle location. In condition 3, the hazards were placed either on the right or left side of the walkway and at variable locations (near, middle or far) (pseudo random order and locations). For all conditions (including when a condition changed), participants were instructed that they “may experience a hazard when walking on this walkway” but not told how, when and where a slip or a trip would occur. They were further instructed to “walk normally in time with the metronome stepping on middle of the black and white tiles”.

**Table 1.** The study protocol.

|  |
| --- |
| *Condition 1: Right leg / fixed location (4 slips and 4 trips)*<br>F1, **<u>N1</u>**, **<u>S1<sub>m</sub></u>**, **<u>T1<sub>m</sub></u>**, F2, **<u>N2</u>**, S2<sub>m</sub>, T2<sub>m</sub>, S3<sub>m</sub>, T3<sub>m</sub>, **<u>S4<sub>m</sub></u>**, **<u>T4<sub>m</sub></u>** |
| *Condition 2: Left leg / fixed location (4 slips and 4 trips)*<br>**<u>S5<sub>m</sub></u>**, **<u>T5<sub>m</sub></u>**, **<u>N3</u>**, S6<sub>m</sub>, T6<sub>m</sub>, S7<sub>m</sub>, **<u>N4</u>**, T7<sub>m</sub>, **<u>S8<sub>m</sub></u>**, **<u>T8<sub>m</sub></u>** |
| *Condition 3: Random leg / random location (4 slips and 4 trips)*<br>**<u>S9<sub>f/R</sub></u>**, **<u>T9<sub>n/R</sub></u>**, T10<sub>f/R</sub>, **<u>N5</u>**, S10<sub>m/L</sub>, S11<sub>n/R</sub>, **<u>N6</u>**, T11<sub>m/L</sub>, **<u>S12<sub>f/R</sub></u>**, **<u>T12<sub>n/R</sub></u>** |

F: free walk (no regulation), N: non-perturbed walk (regulated to usual gait pace), L: left leg, R: right leg, S: slip, T: trip. Locations of the hazards on the walkway (n: near, m: middle, f: far). Gait speed (step length and cadence) was regulated using stepping tiles and metronome adjusted to the participant's usual gait. All participants underwent the same protocol. The bolded letters signify the trials used to in statistical analyses. The purpose and analyses of the F1 and F2 are described in S2 Appendix.

### Measurements

An 8-camera Vicon Bonita motion capture system (Vicon Motion Systems Ltd., Oxford, UK) sampling at 100 Hz was used to collect kinematic data. Thirty four 14-mm diameter reflective markers were attached to the head, trunk, upper and lower limbs according to the Vicon Plug-in-Gait model marker set with a single sacral marker [20]. To avoid loss of important information in for the study purpose (i.e. smoothing out the spikes due to sudden trips and slips), we did not filter the kinematic data. Instead, the quality of the kinematic data and absence of noise in both individual markers and model outputs have been visually confirmed prior to commencement of statistical analyses. Centre of mass (CoM) position was calculated using the Vicon Nexus 1.8.3 full body Plug-in-Gait model. The kinematic variables were calculated from the 3D marker trajectories using custom software developed in MATLAB R2010a (The MathWorks, Inc., Massachusetts, USA).

### Falls and kinematic parameters

A fall was defined by a harness supported load >30% of a participant’s body weight [21]. To examine predictive gait alteration and balance recovery response, the following kinematic parameters were calculated during the step before and after perturbation-onset. As a primary outcome, margin of stability (MoS), a measure of dynamic stability [22], was calculated at step touchdown. The MoS is the anterior-posterior distance (cm) between the closest edge of the base of support (BoS) and the velocity- corrected (extrapolated) sagittal plane centre of mass (XCoM) position. XCoM was calculated as the position of (the vertical projection of) the CoM plus its velocity times a factor 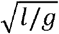 (1 = leg length and g = acceleration of gravity) on the basis of a simple inverted pendulum model [22]. The post-perturbation step MoS was the primary outcome to examine improved balance recovery response. To quantify the magnitude of the balance perturbation caused by the trip/slip, XCoM displacement (cm), the distance between XCoM and the ankle joint of the supporting limb was calculated at the frame prior to touchdown of the leading foot. Step length (cm) relative to the previous step was calculated from foot markers. The XCoM displacement and step length which determine the MoS (i.e. the CoM and BoS states) were two secondary outcomes. In addition, average slip speed (cm/s) was calculated as balance recovery response from slips. The slip initiation (foot strike to the slipping tile) and the slip end (foot lift off from the slipping tile) were visually determined in the Vicon 3D workspace. As measures of predictive gait alterations (i.e. after-effect), during no-perturbation trials, gait speed (m/s), cadence (step/min), step length (cm), toe height (cm) at mid-swing and foot-contact angle relative to the ground (deg) were also calculated and averaged across 4 steps.

### Strategies for balance recovery

Recovery strategies from a trip were classified as (1) lowering strategy: the obstructed limb was quickly lowered to the ground before the obstacle, (2) elevating-contact strategy: the obstructed limb cleared the obstacle after obstacle-contact, and (3) elevating-cross strategy: the foot crossed over the obstacle without contact [23, 24]. Strategies for recovery from a slip were classified as (1) backward and (2) forward stepping when the first recovery foot landed posterior or anterior to the slipping foot, respectively [5, 9].

### Statistical analysis

To test our hypotheses, the regulated non-perturbed walks (N1-N6), the first and last slip (S1, S4, S5, S8, S9 and S12) and first and last trip (T1, T4, T5, T8, T9 and T12) trials in each condition were used in the following planned analyses. Predictive gait alteration was first assessed by examining whether changes existed in gait parameters during regulated non-perturbed walk trials (N1 vs N2), and secondly, change in previous-step kinematics during slip/trip trials at the fixed hazard locations (S1 vs S4 and T1 vs T4). If significant changes (i.e. predictive gait alterations) were observed, transfer between limbs and locations was assessed by investigating whether the changes persisted when a condition was changed without notice (inter-limb transfer: N1 vs N3, S1 vs S5 and T1 vs T5, inter-location transfer: N1 vs N5, S1 vs S9 and T1 vs T9). To confirm the absence of predictive gait alterations at the end of condition 3, kinematics between the first and last non-perturbed walks (N1 vs N6), and previous-step kinematics between the first and last trials for slips (S1 vs S12) and trips (T1 vs T12) were compared. Improvements in balance recovery responses were assessed by examining whether changes existed in recovery-step kinematics at the completion of the condition using random hazard locations (S1 vs S12 and T1 vs T12). Paired *t*-tests were applied for the above comparisons. Changes in the balance recovery strategy (i.e. effects of trial number on the proportion of strategies) were examined by applying the generalized linear mixed model (multinomial or binomial logistic regression) for slip and trip trials separately. IBM SPSS Statistics version 24 (IBM Corp., New York, USA) was used for the analyses. *P*<0.05 was considered statistically significant.

## Results

### Demographics, gait characteristics and dropout

Ten participants (5 female and 5 male) completed the protocol and were included in the analyses. The demographics and usual gait characteristics of the participants (mean ± standard deviation) were as follows: age 29.1 ± 5.6years, height 176.4 ± 11.2cm, weight 70.4 ± 14.8kg, cadence 110.3 ± 7.6step/min, step length 74.3 ± 8.2cm and gait speed 137.3 ± 18.6cm/s. The dominant leg was right for 9 participants.

### Falls

One participant (10%) fell at S1 and 2 participants (20%) fell at S5 (i.e. recorded ≥30% body weight on the harness following the slip). No participants fell in any of the trip trials.

### Gait patterns during no-perturbation trials

Throughout the regulated non-perturbed trials (N1 vs N2 and N1 vs N6), participants walked with similar gait speed, step length, cadence and foot-contact angle (p>0.05) (Fig 2). A significant increase of toe height was observed during the first condition (N1 vs N2, p=0.009). This increased toe height were maintained from right and left legs (N1 to N3, p=0.001) and from fixed to random locations (N1 to N5, p=0.003).

**Fig 2.**
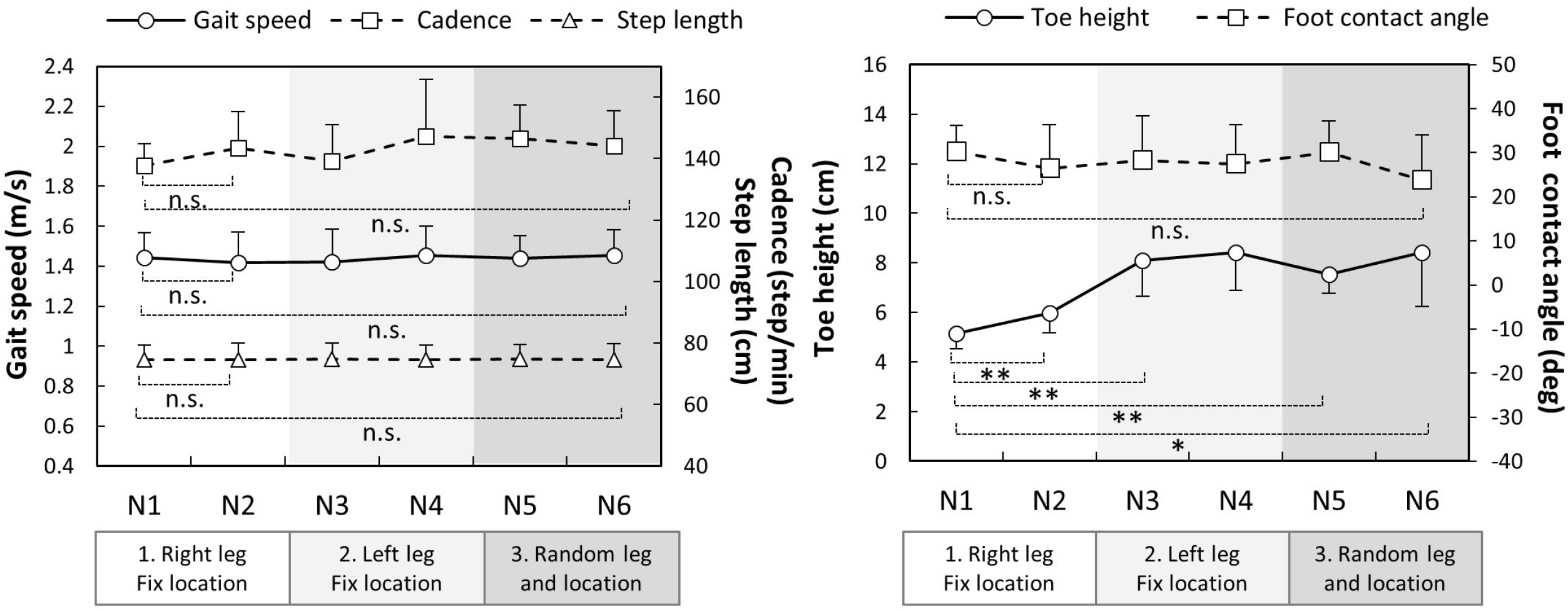
Gait parameters during non-perturbed walk trials (N1 to N6, n=10) interspersed within slip and trip trials throughout the protocol. The dots and error bars are means and standard errors, respectively. * *p* < 0.05, ** *p* < 0.01, n.s. *p* > 0.05

### Kinematics during slip trials

During the first condition (right leg, fixed location), significant changes to previous step kinematics included a greater anterior XCoM displacement (S1 vs S4, *p*=0.009) and reduced MoS (S1 vs S4, *p*=0.002) suggesting predictive gait alteration (Fig 3). Although, these changes were maintained from right to left legs (S1 vs S5, *p*<0.004), they did not maintain in random hazard location (S1 vs S9, *p*>0.096). At the end of the random condition, there were no significant changes in previous step kinematics (S1 vs S12, *p*>0.05) indicating the absence of predictive gait alterations against slips.

**Fig 3.**
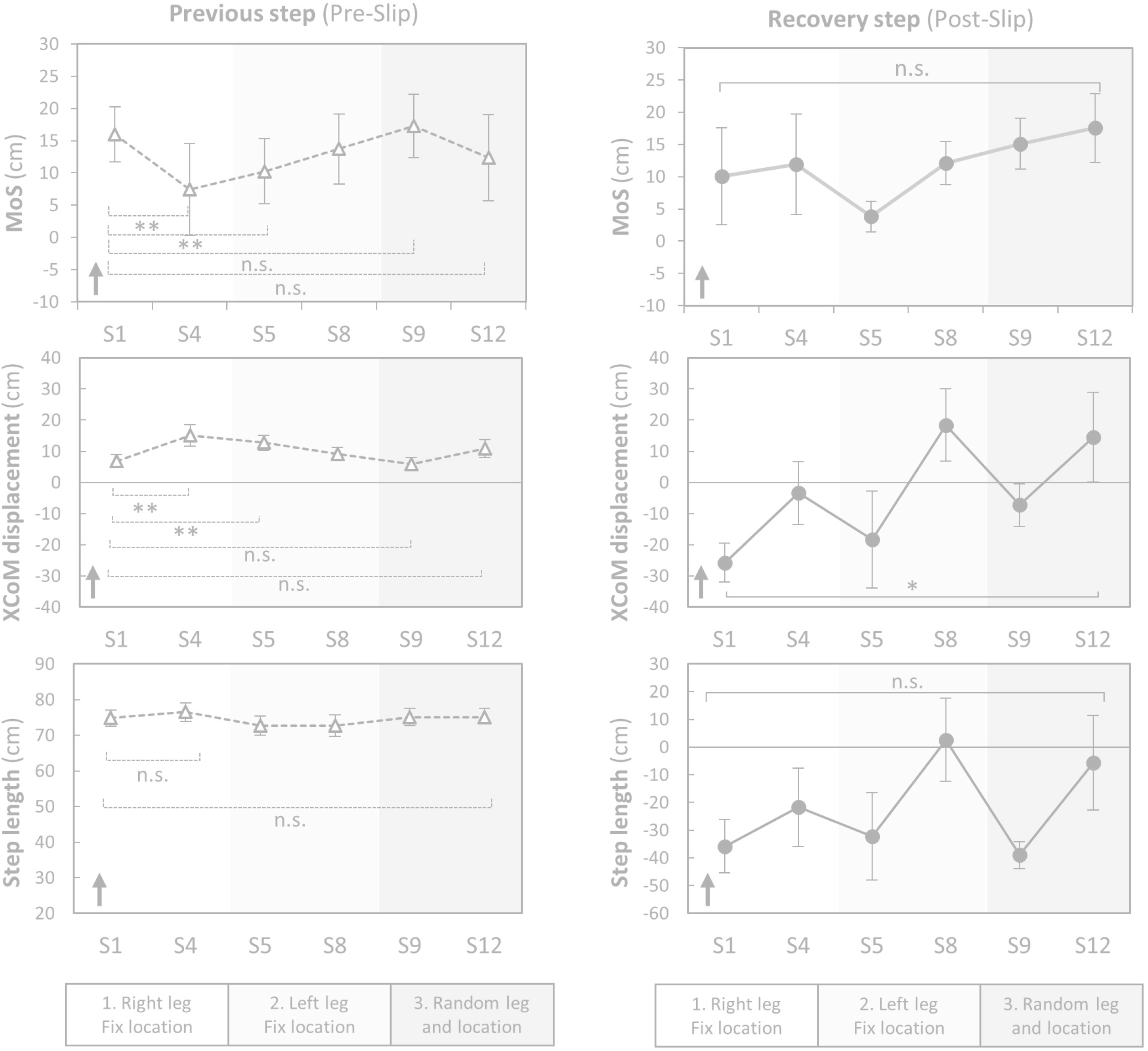
Changes in previous- and recovery-step kinematics during slip trials (n=10). Margin of stability (MoS) was the distance between an extrapolated (i.e. velocity-corrected) centre of mass (XCoM) to the closest base of support limit at foot touch down. XcoM displacement was the distance between the XCoM to the ankle joint of the supporting limb in the sagittal plane. The dots and error bars are means and standard errors, respectively. The arrows indicate the possible directions relating to better stability. S: slip. * *p* < 0.05, ** *p* < 0.01, *** *p* < 0.001, n.s. *p* > 0.05

During a recovery step, significantly less posterior XCoM displacement were observed for the last slip, compared to the first (S1 vs S12, *p*<0.05) suggesting improvements in balance recovery responses (Fig 3). No significant changes in MoS and step length were seen across conditions (S1 vs S12, *p*>0.05). Slip speed also significantly decreased from S1 (135.8 ± 10.4 cm/s) to S12 (82.8 ± 25.9 cm/s, *p*=0.005).

### Kinematics during trip trials

During a step prior to a trip, MoS, XCoM displacement and step length showed no significant changes during all conditions indicating no predictive gait changes (T1 vs T4 and T1 vs T12, *p*>0.05) (Fig 4).

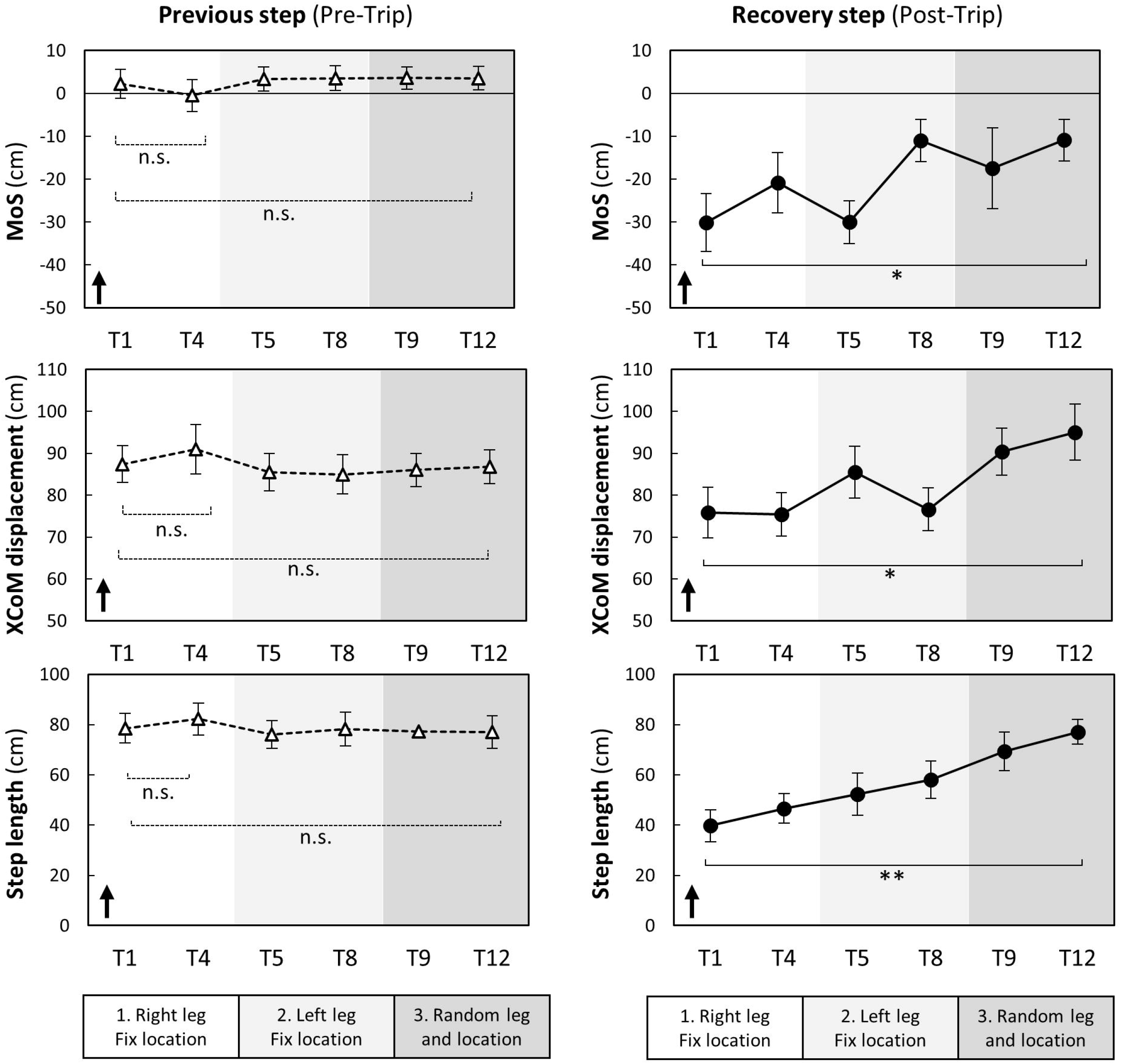

During a recovery step, there was a significant increase in MoS, XCoM displacement and step length between the first and last trip (T1 vs T12, *p*<0.05) suggesting improved reactions to trips (Fig 4).

### Balance recovery strategy

Fig 5 shows stepping strategies taken to recover balance following slips and trips. Backward stepping was most prevalent at the first slip (S1: 90%), which declined to 50% at S8 and increased to 100% at S9 (first slip at random location) and decreased again to 50% at S12. These changes in the proportion of strategies for balance recovery from slips were not significant (*p*>0.05). However, there was a significant change in the proportion of strategies for balance recovery from trips (*p*<0.01). As shown in Fig 5, the lowering-contact strategy was most prevalent at the first trip (T1: 90%), decreasing to 10% at T12. In turn, the proportions of the elevating-contact strategy (T1: 0% to T12: 50%) and elevating-cross strategy (T1: 10% to T12: 40%) increased across the trials.

**Fig 5.**
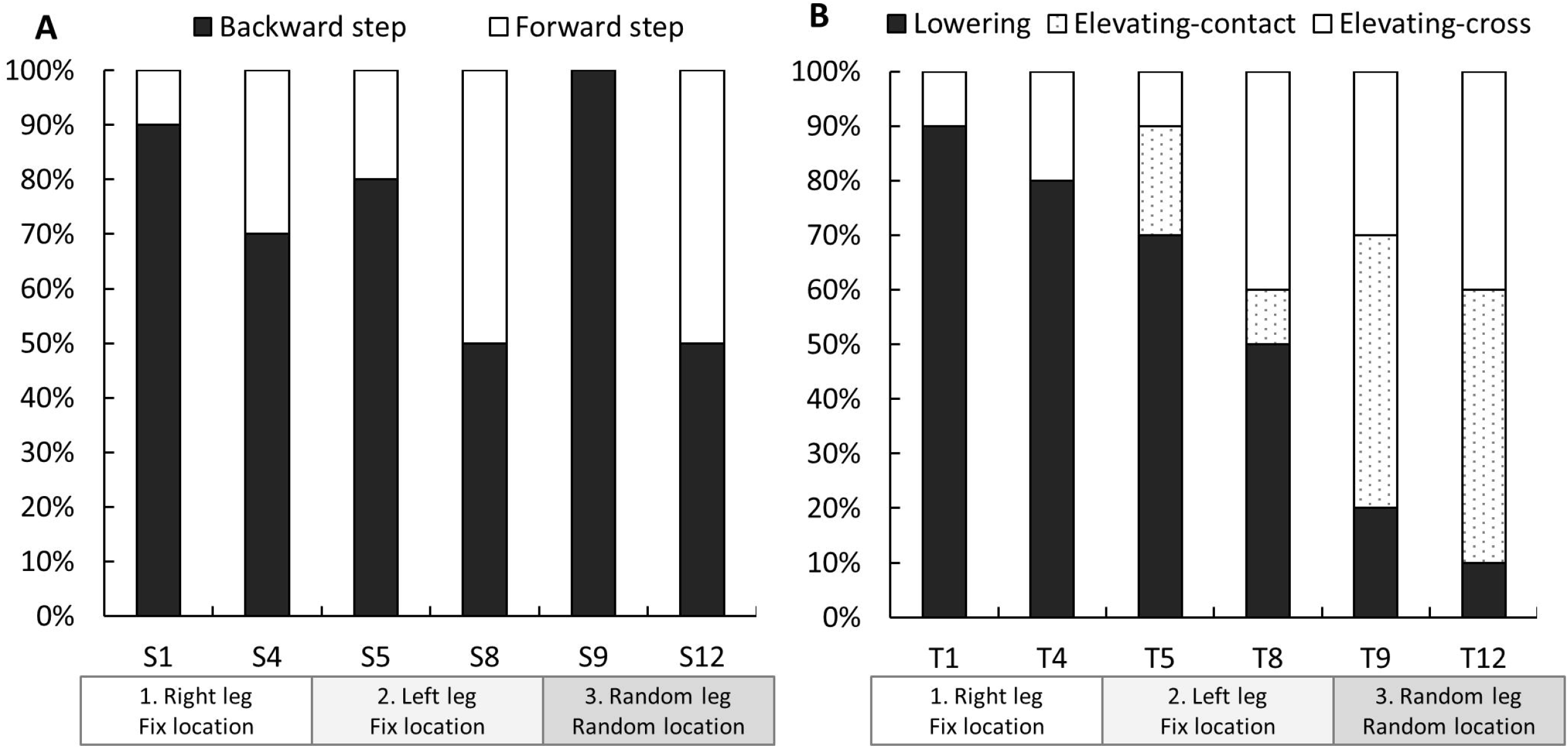
Strategies for balance recovery after slips (A) and trips (B) (n=10). S: slip, T: trip. Changes in proportions of balance recovery strategies used were examined by applying the generalized linear mixed model (multinomial or binomial logistic regression). A significant effect of trials was observed in the trip trials (*p*<0.01). This indicate that the proportions of the lowering strategy in response to trips decreased (T1: 90% to T12: 10%) and the elevating-contact (T1: 0% to T12: 50%) and elevating-cross (T1: 10% to T12: 40%) strategies increased. There was no significant effect of trials in the proportion of strategies during the slip trials.

## Discussion

This study aimed to examine balance recovery responses to repeated trips and slips while minimising predictive gait alterations. By repeatedly delivering trips and slips using a walkway with stepping tiles and a metronome, we could successfully regulate gait speed, step length, cadence and examine changes in reactive balance control. At the completion of training, MoS after a trip was significantly increased indicating improved balance recovery responses to unpredictable trip hazards. However, it was less clear if the training improved balance recovery responses from slips.

### Predictive gait alterations and transfer

Previous studies have shown the most common predictive gait alteration (or feedforward adaptation) to repeated slips is an anterior shift of the CoM; a strategy employed to counteract the effect of backward trunk rotation [5, 9, 15, 25]. In this study, we observed an anterior shift of XCoM (additional analysis confirmed significantly forward CoM position but not velocity) even though trips (a perturbation that propels the CoM forward) were interspersed with slips. Although this shift of CoM transferred from one leg to the other when the slip location was fixed, it did not transfer to slips given at random locations. These findings can explain previous studies that have found training effects are transferrable from a movable platform to a slippery floor [11], short to long slips [12], and from one leg to the other [13] i.e. situations where hazard locations are sufficiently alike to allow the same predictive CoM control strategy. However, our findings showing this anterior shift of CoM strategy did not transfer to unpredictable slip hazard locations when slips and trips were interspersed, suggest it is also unlikely to generalize to unpredictable hazards encountered in everyday life.

An anterior shift of the CoM may pose a risk to gait stability because it is related to reduced pre-slip MoS (Pearson’s correlation coefficient *r*=-0.98) indicating a greater vulnerability to trips. Bhatt et al. also found predictive gait alteration to repeated slips leads to greater forward trunk rotation when a trip is unexpectedly encountered, and the forward shift of CoM position acquired during slip training dissipates after mixed slip and trip training [24]. Therefore, while predictive control of the CoM can reduce perturbation impacts and may assist initial learning of balance recovery skills, mixed training appears to minimize poor responses to novel perturbations with opposite CoM displacements.

Increased toe clearance has previously been reported as a predictive gait alteration to repeated trip and/or slip exposures [16, 24]. We found elevated toe height (+3 cm) during gait transferred from right to left legs and from fixed to random hazard locations and also assisted obstacle clearance in later trials (i.e. employment of elevating strategies). Elevated toe clearance may therefore be a predictive strategy that generalizes to safe negation of unpredictable perturbations encountered in everyday life.

### Balance recovery responses from slips

Our regulated gait protocol precluded most predictive gait alterations previously reported (i.e. shorter step length [14, 24], flat-foot landing [5, 14], anterior shift of CoM [5, 9, 15, 25]. With these regulation, our results show participants reduced slip speed and post-slip XCoM displacement likely due to improvements in balance recovery responses. It is possible that slip speed was decreased by reduced muscle activation onset, quicker peak activation times [26] and co-activation of the lower-extremity muscles [26]. If improvements in the sensorimotor system can be obtained through repeated exposure to unpredictable perturbations, it would likely be generalizable in daily life situations where individuals cannot predict all fall hazards but need immediate and appropriate responses.

Previous studies have reported a 90-100% reduction of backward balance loss from the 1^st^ slip to the 5^th^ trial in response to repeated slips in young adults [5, 9]. In contrast, backward stepping in our study (akin to backward balance loss) reduced by only 40% in the first 8 slips (S1-S8), likely because the interspersed slips and trips did not allow large predictive alterations to occur. Moreover, all participants (100%) took a backward step when a slip was encountered at the new location (S9). These backward steps were most likely related to the negative recovery-step XCoM displacement and step length observed during S9. Unlike predictive gait alterations, that occur quickly when the type and location of hazard have been revealed, improvement in balance recovery responses (that possibly involve motor skill learning and improved muscle activation) probably occur more gradually and require longer-term and higher-dose training than the current protocol.

### Balance recovery responses from trips

We found improved balance recovery responses from trips as indicated by increases in recovery-step MoS during recovery. Following repeated exposure to trips, participants were more likely to clear the obstacle, rather than lower the obstructed limb immediately to the floor. This change in strategy likely contributes to improved stability because the lowering strategy (terminating the step before the obstacle) causes a greater gait disruption and is unlikely to assist in arresting the forward momentum of the COM (instability). The improved strategy type and stability (MoS) observed was possibly due to improved use of hip-flexor muscles and improved support limb push-off, as we have seen these to be important for successful trip recovery in previous studies [17, 18]. Pijnappels et al. reported balance recovery from a trip requires support limb power to propel the CoM upwards and provide time and space for an adequate recovery step to arrest the angular momentum of the COM [17, 18]. Faster initiation of the recovery step [27] may also constitute a crucial component of successful balance recovery.

### Study limitations

We acknowledge certain limitations of this study. First, although the protocol aimed at precluding predictive gait alterations and was successful in keeping most gait parameters similar to normal gait, participants walked with an elevated toe clearance throughout the trials and this would have contributed to the improved reactions to trips observed. Second, the manual control of the switch to trigger the trip hazard yielded some variability in timing, such that the trip would not have always been encountered at precisely mid-swing. Third, the use of a metronome and stepping tiles were necessary to maintain the normal gait speed and ensure consistent perturbations, they may also have diminished the natural variability during gait and increased the attentional demands of gait. Fourth, our study did not examine retention effects, which require further investigation to appreciate the utility of perturbation training for fall prevention in daily life. Fifth, our small study sample did not allow correction for multiple comparisons (e.g. Bonferroni), therefore, may be subjected increased type one error. Finally, the generalizability of the study findings to older adults needs to be examined in future studies.

## Conclusions

Even in the absence of most predictive gait alterations, balance recovery responses to trips and to a lesser extent to slips were improved through exposure to repeated unpredictable perturbations. With the exception of elevated toe clearance, most predictive gait alterations to perturbations at a fixed location is likely not generalizable in daily life situations where perturbation type and location could not be predicted. A common predictive gait alteration to lean forward immediately before a slip was not useful when the perturbation location was unpredictable. The findings suggest that training of balance recovery responses to unpredictable perturbations may be beneficial to fall avoidance in everyday life. Further studies are needed to examine retention, kinetic and physiological mechanisms, and generalizability of our findings to older adults at risk of falling.

## Acknowledgements

We express our deep gratitude all the participants of this study.

## Supporting information

**S1 Dataset. Dataset used in this study.**

**S2 Appendix. An analysis on free gait without gait regulation.**

